# A diagnostic primer pair to distinguish between *w*Mel and *w*AlbB *Wolbachia* infection - A simplified real-time PCR approach for use in field collected *Aedes aegypti* in dengue reduction trials

**DOI:** 10.1101/2021.06.03.446891

**Authors:** Meng-Jia Lau, Ary A. Hoffmann, Nancy M. Endersby-Harshman

## Abstract

Detection of the *Wolbachia* endosymbiont in *Aedes aegypti* mosquitoes through real-time PCR assays is widely used during and after *Wolbachia* releases in dengue reduction trials involving the *w*Mel and *w*AlbB strains. However, primers applied in current successful *Wolbachia* releases cannot be used in a single assay to distinguish between these strains. Here, we developed a new diagnostic primer pair *wMA* which can detect *w*Mel or *w*AlbB infection in the same assay. We also tested current *Wolbachia* primers and show that there is variation in their performance when they are used to assess the relative density of *Wolbachia*. The new *wMA* primers provide an accurate and efficient estimate of the presence and density of both *Wolbachia* infections.

## Introduction

The bacterium, *Wolbachia*, is providing an increasingly popular method to inhibit dengue virus transmission in the mosquito, *Aedes aegypti. Wolbachia-infected* populations involving the *w*Mel strain have now been successfully established in *Ae. aegypti* in regions including northern Australia, Brazil and Indonesia [1–3], while *w*AlbB-infected *Ae. aegypti* have been established in Malaysia [4]. Detection of the *Wolbachia* endosymbiont in *Ae. aegypti* mosquitoes is a standard requirement for good laboratory practice during *Wolbachia* mosquito releases in dengue reduction programs and for tracking *Wolbachia* invasions in the field [4, 5]. Real-time PCR and High Resolution Melt (HRM) assays (SYBR^®^ equivalent/non-probe) have been developed that enable detection and *Wolbachia* density estimation for the strain of interest [6–8]. However, difficulties can arise in using these assays when there is a need to detect *Wolbachia* and distinguish between multiple *Wolbachia* strains. In experiments where superinfected lines are used [9], or where mosquitoes carrying different single infections need to be distinguished for experiments or in field collected samples [10], different assays are currently required. Given that both *w*Mel and *w*AlbB strains are now actively being used in field releases and that each strain may have advantages in particular situations, the requirement for multiple strain identification is likely to increase in the foreseeable future.

In previous work, we have used a *Wolbachia-specific* primer pair, w1 [7], which targets a conserved locus VNTR-141 containing tandem repeats [11]. This pair of primers works efficiently in amplifying *w*Mel and *w*MelPop infections in a real-time PCR and HRM assay, but achieves only poor amplification of *w*AlbB [10]. As well as being used for *Wolbachia* detection, primers are needed for quantification of *Wolbachia* density in mosquitoes. There are various *Wolbachia* specific primers for *w*Mel, *w*AlbB or *w*MelPop [9, 12–14], but currently there is no standardised assay for *Wolbachia* screening that is comparable between strains and that can be used to compare results between laboratories. Although cross-laboratory comparability may not be a realistic aim when using a SYBR^®^ equivalent/non-probe-based assay, the use of extra internal controls can make these assays robust for relative density estimates, improving consistency within laboratory experiments [7, 10].

In this study, we developed a diagnostic primer pair that can detect and distinguish between the *w*Mel and *w*AlbB infections and also provides an estimate of *Wolbachia* density. In addition, we assessed primer efficiency of some other published primers for *Wolbachia* in mosquitoes and looked for cross-amplification between strains.

## Materials and methods

To develop the new primers, we screened for differences between the *w*AlbB and *w*Mel strains and then focussed on the sequences of gene WD_RS06155 in *w*Mel and its analogue in *w*AlbB. We then developed a new pair of primers designated *wMA* (Table 1) to distinguish *Wolbachia w*Mel and *w*AlbB in a single run of a qPCR assay, based on two base-pair mismatches at the 3’- end of each primer, which resulted in the Tm peak for *w*AlbB being separated from that of *w*Mel.

**Table 1.**
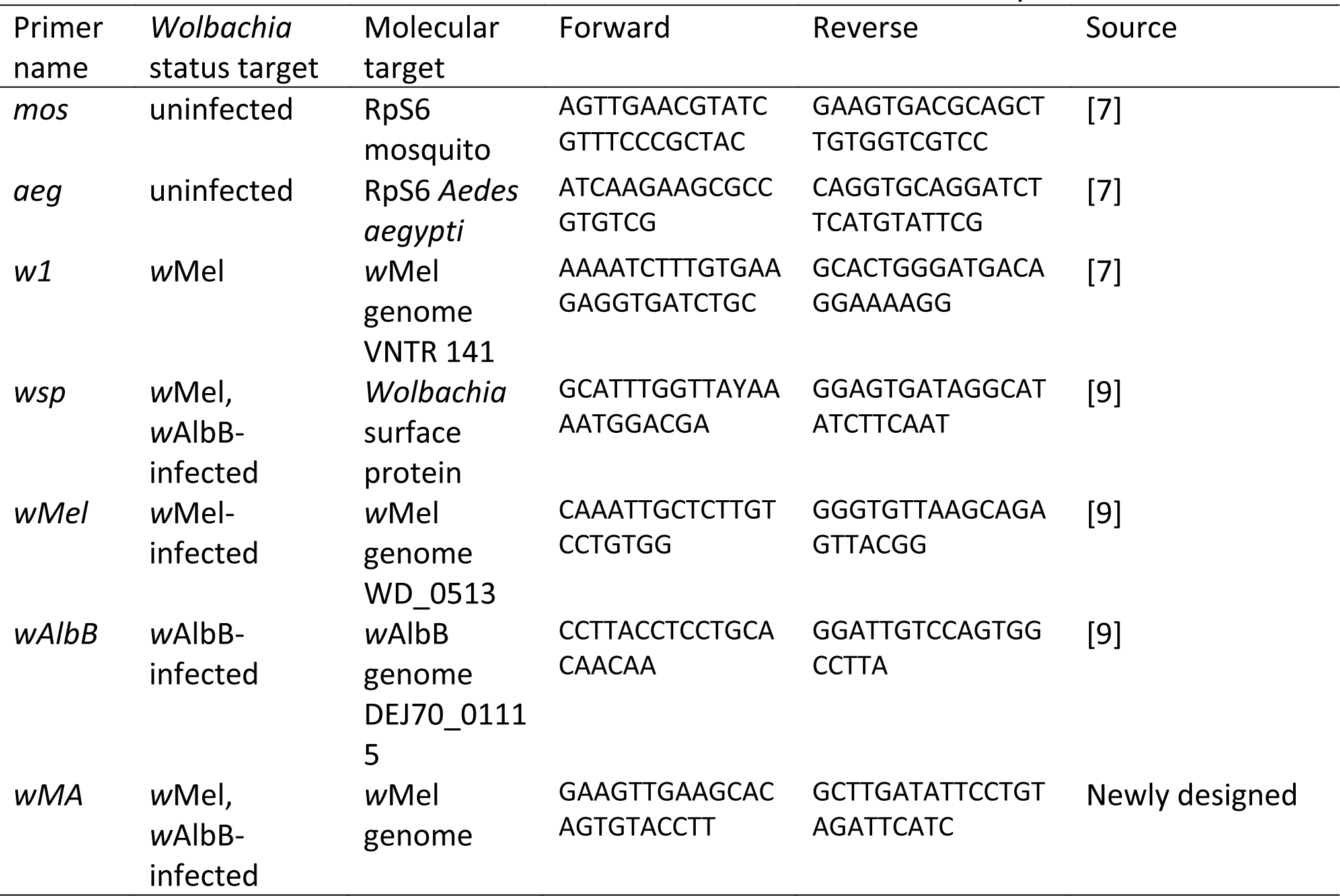
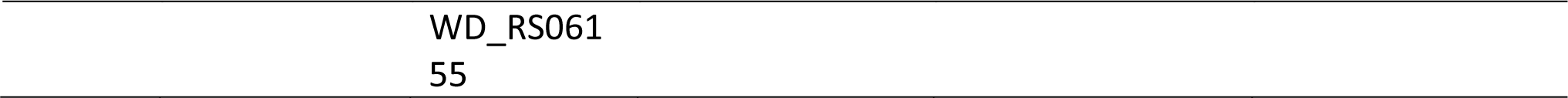
Primers for detection of *Wolbachia* strains and estimation of density

### Sample preparation and LightCycler^®^ efficiency test

The *w*Mel and *w*AlbB-infected *Ae. aegypti* were tested against lines transinfected previously [15, 16]. The *w*Mel line was collected from Cairns, Australia in 2019 from regions that had been invaded several years earlier [2, 12], while the *w*AlbB line was derived from a *w*AlbB infected line crossed to an Australian background and maintained in the laboratory.[13]. An uninfected line which was developed from *w*Mel-infected line after tetracycline treatment was also used in this study.

Mosquitoes were reared with TetraMin^®^ fish food tablets in reverse osmosis (RO) water until the adult stage [17]. We used a mixture of young (4±1 days since eclosion) and old (38 ±1 days since eclosion) mosquitoes from *w*Mel-infected [16], *w*AlbB-infected [15] and uninfected strains that were killed in absolute ethanol before Chelex^®^ DNA extraction. In the standard procedure, DNA was extracted in 250 μL 5% Chelex^®^ 100 Resin (Bio-Rad Laboratories, Hercules, CA) and 3 μL of Proteinase K (20 mg/ mL, Bioline Australia Pty Ltd, Alexandria NSW, Australia). The DNA concentration in the Chelex^®^ 100 Resin solution was measured using a Quantitation Qubit^Tm^ 1X dsDNA HS Assay Kit, and then diluted ten times before making a three-fold dilution series to test the efficiency of currently-used *Wolbachia* primers in a real-time PCR assay. We also diluted the solution six times before making a three-fold dilution series to investigate the influence of DNA and Chelex^®^ concentration.

For the real-time PCR and HRM, we used a LightCycler^®^ 480 High Resolution Melting Master (HRMM) kit (Roche; Cat. No. 04909631001, Roche Diagnostics Australia Pty. Ltd., Castle Hill New South Wales, Australia) and IMMOLASE™ DNA polymerase (5 U/μl) (Bioline; Cat. No. BIO-21047) as described by Lee et al. (2012). The PCR conditions for DNA amplification began with a 10 minute pre-incubation at 95°C (Ramp Rate = 4.8 °C/s), followed by 40 cycles of 95 °C for 5 seconds (Ramp Rate = 4.8°C/s), 53°C for 15 seconds (Ramp Rate = 2.5°C/s), and 72°C for 30 seconds (Ramp Rate = 4.8°C/s).

Three technical replicates were run for each sample of each dilution and a graph was plotted showing the log3 [dilution factor] (x-axis) against mean Cp (y-axis) and a linear trend line (y = mx + c) was fitted. Slope (m) and R^2^ values were recorded so that PCR amplification efficiency (E) could be evaluated with the equation:

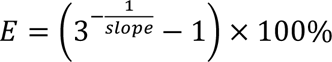

### Primer crossing points comparison and density estimation

Following the efficiency study, we tested for Cp value differences between primers for different *Wolbachia* strains to assess suitability for relative *Wolbachia* density estimation from a total of 16 *Wolbachia-infected* mosquito samples, which were extracted using Chelex^®^ resin and then diluted ten times before real-time PCR.

## Results and Discussion

### Diagnostic primer design

In this study, we developed a diagnostic primer pair that can detect and distinguish between the *w*Mel and *w*AlbB infections in *Ae. aegypti*, which is important in simplifying current approaches for *Wolbachia* identification. The newly designed primer pair *wMA* has distinctive Tm values between *Wolbachia w*Mel (82.6 ± 0.03 °C) and *w*AlbB (80.4 ± 0.02 °C) screening (Figure 1c). The high resolution melt produces two joined peaks when the template contains both *Wolbachia w*Mel and *w*AlbB DNA (Figure 1d).

**Figure 1.**
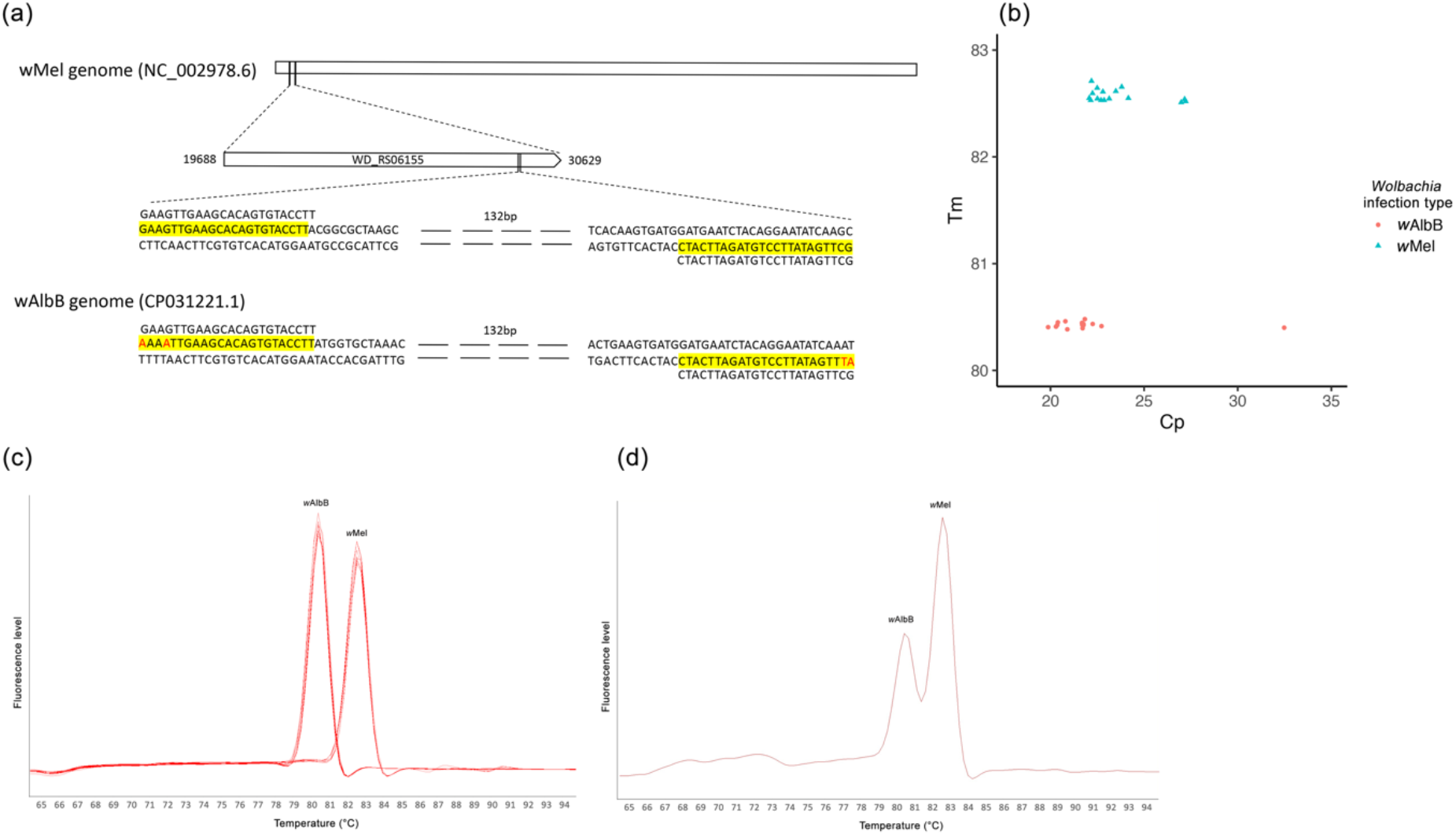
Development of primers to detect *Wolbachia w*Mel and *w*AlbB infection in *Ae. aegypti*. (a) The new primer pair *wMA* aligns to a region in gene WD_RS06155 of *w*Mel, and aligns to its analogue in *w*AlbB genome which had two base-pair mismatches at the 3’- end; (b) the *wMA* primers showed distinct Tm values for *Wolbachia w*Mel (82.6 ± 0.03 °C) and *w*AlbB (80.4 ± 0.02 °C), the x axis represents the crossing point and the y axis represents the amplicon melting temperature; (c) the *wMA* primers showed distinct Tm values for *Wolbachia w*Mel and *w*AlbB; (d) the *wMA* primers showed two Tm peaks when mixing DNA template of *w*Mel and *w*AlbB-infected *Ae. aegypti*.

### Primer efficiency test and Cp values comparisons

We tested the efficiency of each of the primers used for screening *Wolbachia* in *Ae. aegypti* (Table 2) by using a threefold dilution series. The efficiencies of all primers were within the acceptable range (85% −110%, Table 2, Figure 2) [18] and the efficiency curves all showed an R^2^ valued greater than 0.99.

**Table 2.** Primer efficiency for primers used in detection of *Wolbachia* strains and estimation of density. DNA is extracted in 250 μL 5% Chelex^®^ 100 Resin and then diluted ten times before making a three-fold dilution series.

| Colony | Primers | Slope of graph | $R^2$ | Efficiency | DNA concentration in Chelex<sup>®</sup> 100 Resin (ng/μL) |
| --- | --- | --- | --- | --- | --- |
| Uninfected | *mos* | -1.566 | 0.999 | 101.659% | 5.12 |
| Uninfected | *aeg* | -1.531 | 0.999 | 104.946% | 5.12 |
| wMel | *w1* | -1.644 | 0.999 | 95.109% | 5.24 |
| wMel | *wM* | -1.677 | 0.998 | 92.559% | 5.24 |
| wMel | *wsp* | -1.767 | 0.999 | 86.195% | 5.24 |
| wMel | *MA* | -1.769 | 0.999 | 86.107% | 5.24 |
| wAlbB | *MA* | -1.730 | 0.997 | 88.732% | 6.87 |
| wAlbB | *wsp* | -1.755 | 0.999 | 87.032% | 6.87 |
| wAlbB | *wA* | -1.741 | 0.993 | 87.953% | 6.87 |

**Figure 2.**
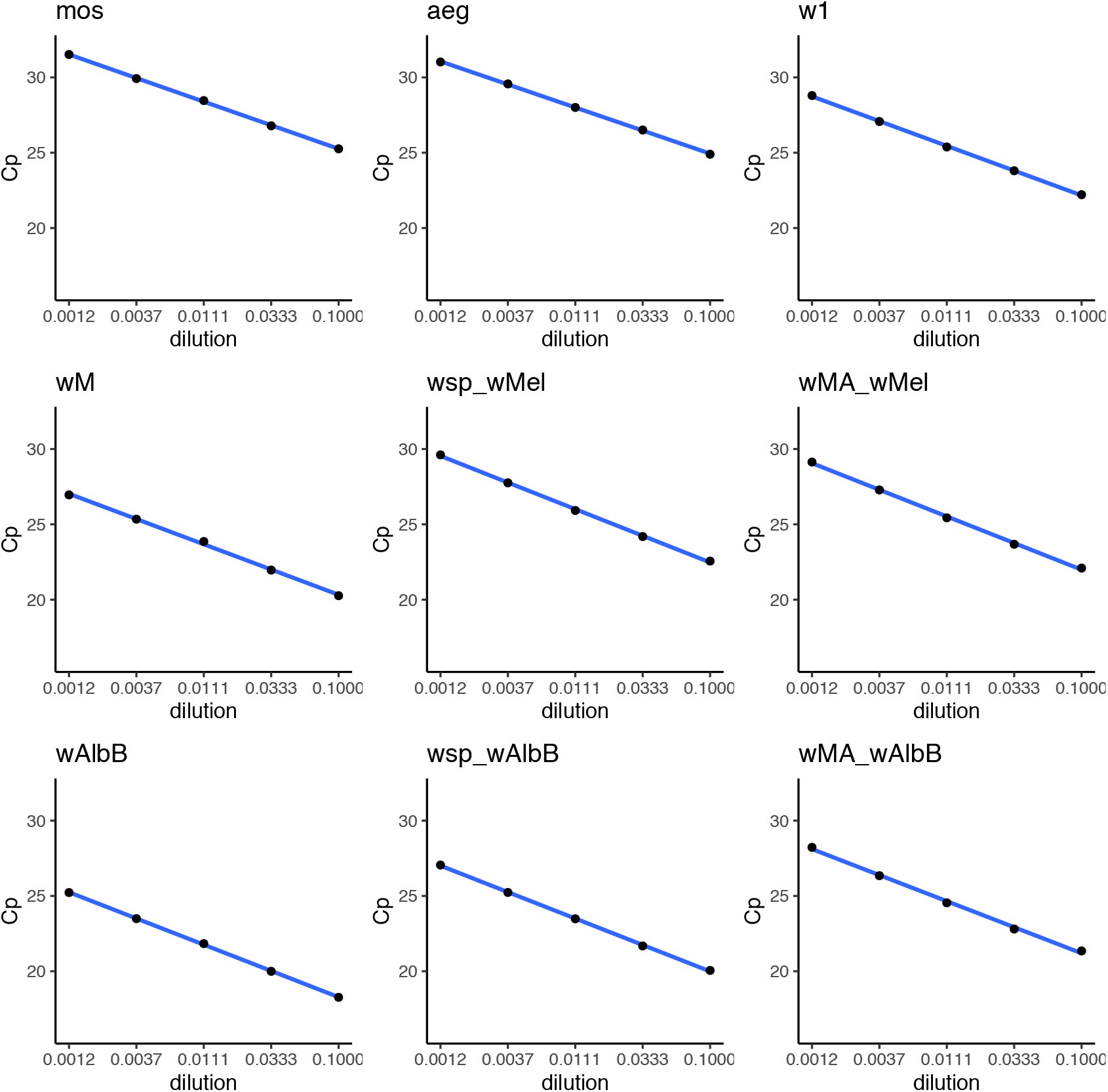
Primer efficiency for detection of *Wolbachia* strains and estimation of density. DNA is extracted in 250 μL 5% Chelex^®^ 100 Resin and then diluted ten times before making a three-fold dilution series. The primer names are defined in Table 2.

However, we found the amplification curve increase showed inhibition at the first dilution (Figure 3) for each of the primers. We also found the same phenomenon when DNA was first diluted six times instead of ten times which resulted in outliers occurring in efficiency curves (Sup. Figure 1, 2). These results highlight a potential risk of lowering the relative density estimate in *Wolbachia* screening when using highly concentrated Chelex^®^-extracted DNA solution.

**Figure 3.**
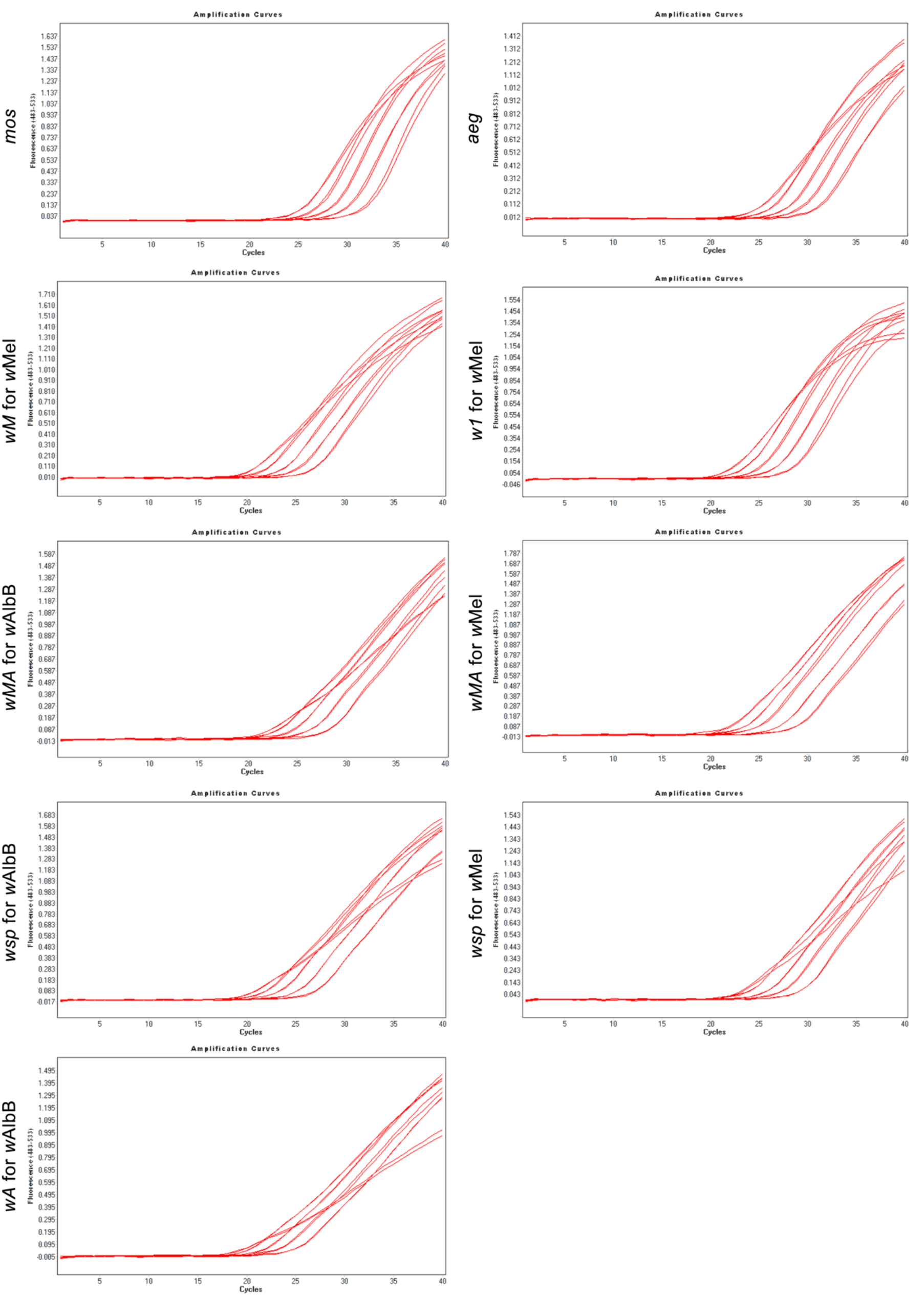
Variation in the shape of the PCR amplification curves when Cp values are fitted with the standard curve. The curves from left to right represent amplification curves of 1/10, 1/30, 1/90, 1/270 and 1/810 DNA dilution from 250 μL 5% Chelex^®^ 100 Resin. The primers are defined in Table 2.

We noticed that primers had different Cp values even when screening the same individual organism (*Ae. aegypti, w*Mel or *w*AlbB) using the same DNA concentration, even though these primers all have a high efficiency. We therefore tested the Cp ranges of the primers and confirmed variation when using them (Figure 3), which would be expected to result in differences in relative density estimates. The relationship between Cp values of different primers all fit into a linear relationship, with R^2^ greater than 0.97, whereas the coefficient varies from 0.83 to 1.05 (Fig. 4).

**Figure 4.**
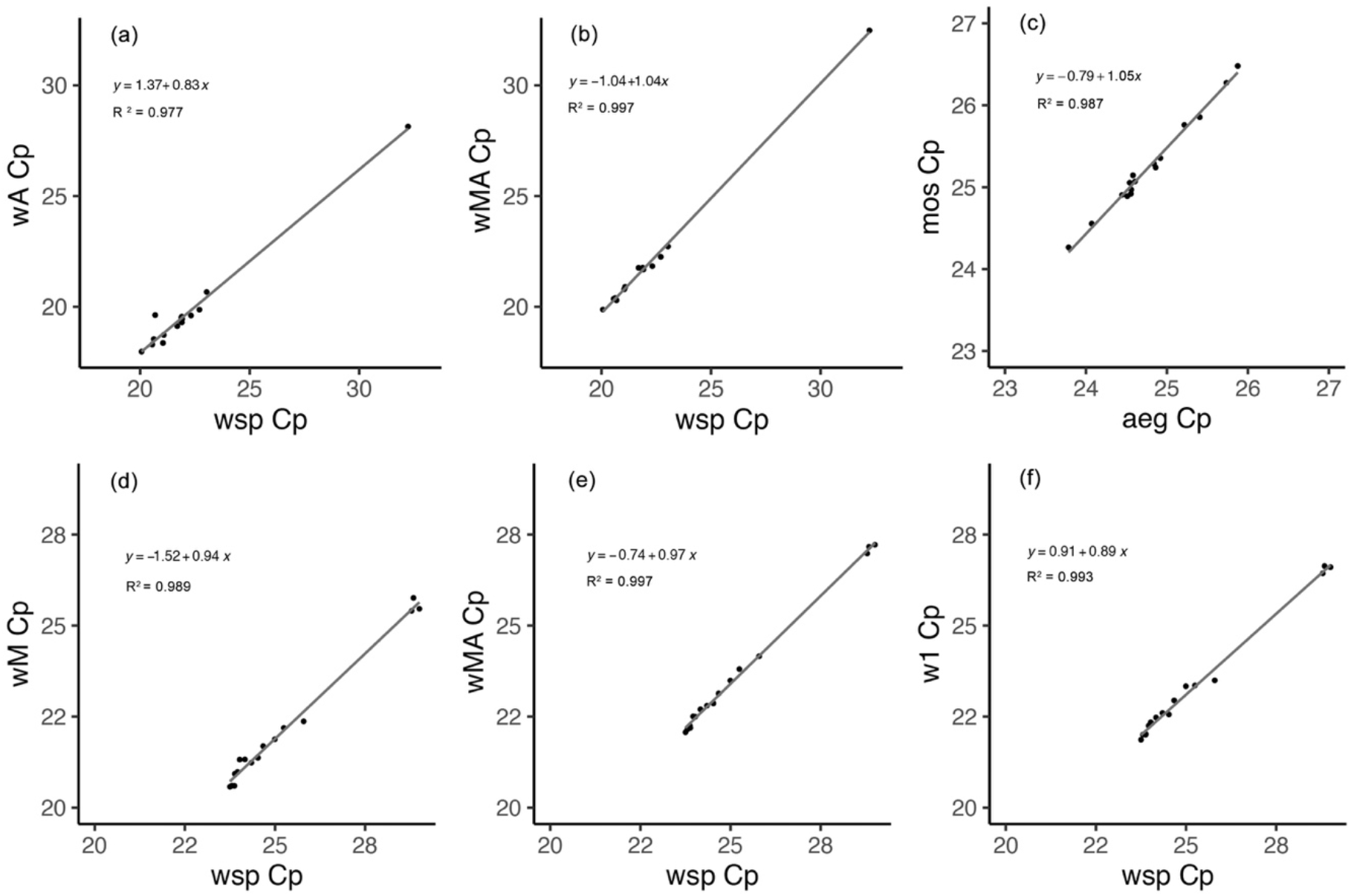
Variation in Cp values when using different *Wolbachia* primers for the same samples. (a) *wA* and *wsp* primers in *Wolbachia w*AlbB screening; (b) *wMA* and *wsp* primers in *Wolbachia w*AlbB screening; (c) *mos* and *aeg* primers; (d) *wM* and *wsp* primers in *Wolbachia w*Mel screening; (e) *wMA* and *wsp* primers in *Wolbachia w*Mel screening; (f) *w1* and *wsp* primers in *Wolbachia w*Mel screening.

This finding could not be explained fully by pipetting error and PCR inhibition [19, 20] as PCR efficiency was always high. Inhibition effects on the DNA amplification can vary not only when using different primers, but also when the DNA concentration varies. The intercepts of these Cp values differ from −1.52 to 1.37, indicating that there may be different copies of genes inside mosquito cells [21, 22], which can be variable under different environmental conditions [23, 24]. As a result, care is needed when choosing primers for assessing the relative concentration of *Wolbachia*

## Conclusions

The primers provide a useful approach for detecting both of the currently released *Wolbachia* infections. Other options like multiplex probe assays and using DNA extraction kits to purify DNA are likely to cost more, and the Chelex^®^ DNA extraction remains an easy and economic method for high-throughput *Wolbachia* screening [4]. Given potential variation within infections, we would still recommend undertaking comparisons between *Wolbachia* primers described in our study and other *Wolbachia-specific* primers in different field contexts. Comparisons with universal *Wolbachia* primers should also be undertaken before using the newly-designed primers in *Wolbachia* density calculations. The *wsp* primers still represent a useful pair of universal primers for amplifying the *Wolbachia* surface protein gene which has been used as a *Wolbachia* diagnostic for decades [14]. For the *Wolbachia w*Mel or *w*AlbB infection in *Ae. aegypti*, our newly-designed primer pair, *wMA*, not only identifies *w*Mel and *w*AlbB at the same time, but is also correlated with density estimates based on *wsp*, with correlation estimates for both *w*Mel and *w*AlbB close to 1. Thus, the new primers have the potential to be an accurate and efficient approach for large-scale *Wolbachia* screening.

## Acknowledgements

We thank Perran Stott-Ross and Jason Axford for providing the mosquito samples. We also thank support of Jasper Loftus-Hills award offered by the Faculty of Science, the University of Melbourne. This research is also funded by the National Health and Medical Research Council (1132412, 1118640, www.nhmrc.gov.au) and the Wellcome Trust (108508, wellcome.ac.uk).

